# Buried but unsafe – defoliation depletes the underground storage organ (USO) of the mesic grassland geophyte, *Hypoxis hemerocallidea*

**DOI:** 10.1101/2021.03.18.435941

**Authors:** Craig D. Morris

## Abstract

Mesic grasslands in South Africa (> 650 mm a^-1^ MAP) are rich in herbaceous forbs, which outnumber grass species by more than 5 to 1. Many of these forbs have underground storage units (USOs), such as thickened rootstocks, rhizomes, bulbs, or corms, that provide resources (non-structural carbohydrates, minerals, and water) enabling them to resprout after dry, frosty winters, and fire. However, despite their extensive biomass and reserves ostensibly protected underground, geophytic mesic grassland forbs can be severely depleted or extirpated by chronic trampling and grazing of their aerial parts by livestock. This study examined a possible explanation for forb demise in overgrazed grassland by investigating, in a pot trial, whether the growth of forbs and the size of their USOs are negatively affected by simulated green leaf loss. In a 2×2 factorial (clipped vs. unclipped x spring regrowth in the dark vs. light), five replicate plants of *Hypoxis hemerocallidea*, a common mesic grassland forb that resprouts from a corm, were subject to six severe (clipped to 80 mm) defoliations during the growing season and regrown in spring under full or restricted light to measure stored reserve contribution to regrowth. Defoliated plants were resilient to defoliation during the growing season, matching the total biomass production of unclipped plants, though cutting reduced the number of leaves by ¬60% and flowers by almost 85%. Spring regrowth on stored reserves equalled that from reserves plus concurrent photosynthesis, indicating the value of USOs for regrowth. However, there was a marked carry-over effect of previous season defoliation, resulting in a one-third reduction in shoot growth and 40% fewer inflorescence in spring. Crucially, corm mass was more than halved by clipping. Above-ground spring growth was linearly related to corm mass. It was concluded that buried stored reserves are not protected by recurrent disturbance to aerial plant parts and that continued diminishment of USOs under chronic disturbance by overgrazing or frequent mowing would weaken and likely eventually kill plants, reducing forb species richness. Lenient management by infrequent summer mowing or grazing at moderate stocking rates combined with periodic rotational full season resting and dormant-season burning is recommend to maintain the USOs and vigour of forbs in mesic grassland.

## 1. Introduction

Grasslands, including those in southern Africa (Cowling and Hilton-Taylor, 1997; Bond and Parr, 2010), are among the most speciose plant communities on the planet at point and patch scales (sub-metre to 100 m^2^), despite being subject to frequent and often intense disturbance by fire, grazing, or mowing (Wilson et al., 2012; Martorell et al., 2017). Most of the plant species diversity of mesic grassland in South Africa (> 650 mm a^-1^ MAP) comprises herbaceous monocotyledonous and dicotyledonous herbaceous forbs (hereafter referred to as ‘forbs’), which contribute more than 80% of the species richness (Morris, 2004; Uys et al., 2004). The diverse suite of forbs co-occurring with grasses in South African mesic grassland has coevolved with recurrent dry season (winter to early spring) fires fuelled by senescent grass phytomass accumulated in the growing season (Everson et al., 1988; Bond et al., 2003). Forbs resprout rapidly after a burn from buds on underground storage organs (USOs), such as woody rootstocks, bulbs, or corms (Uys, 2006; Zaloumis and Bond, 2016; Klimešová et al., 2019), that protect growing perennating buds from fire and frost while providing a store for energy for rapid regrowth of leaves and inflorescences in spring (Dafni et al., 1981; Clarke et al., 2013; Pausas et al., 2018; Lubbe et al., 2021). Consequently, mesic grassland forbs can consequently withstand frequent topkill by frost, fire, or mowing (Fynn et al., 2004; Morris et al., 2021). Investment in below-ground structures such as USOs does, however, render forbs vulnerable to soil disturbance: destruction of damage to USOs resulting from cultivation for cropping or afforestation can eliminate or severely reduce populations of grassland forbs (Bond and Zaloumis, 2016; Silveira et al., 2020).

Another type of disturbance inimical to mesic grassland forbs is overgrazing by livestock. Light to moderate levels of grazing can sustain forb populations (Joubert et al., 2017; Parrish, 2017) but heavy grazing, especially if relentless, can reduce plant species diversity and markedly transform species composition (O’Connor, 2005; Uys, 2006; Chamane et al., 2017a). Scott-Shaw and Morris (2015) reported that intense grazing can deplete up to 84% of the species richness of indigenous forbs. Frequent summer mowing can also be detrimental to certain forb species (Fynn et al., 2005; Valkó et al., 2012). The process whereby grazing animals and mowing can negatively impact mesic forb plants and eventually reduce their populations is not clear as the regenerative organs and energy supplies of such forbs are underground, out of the reach of the mouths and hooves of herbivores and the blades of mowers (Cole, 1995): only 20% of grassland biomass is above ground (Ottaviani et al., 2020). This study examines a possible indirect mechanism to explain the demise of forbs under grazing by testing whether repeated disturbance to the above-ground parts of a widespread mesic forb, *Hypoxis hemerocallidea* Fisch., C.A.Mey. & Avé- Lall., has an impact on the size of its underground storage organ (a corm) and its ability to support resprouting in the following spring.

Only recently has the extent and degree of above-ground damage to forbs by grazing animals been quantified. Chamane et al. (2017b) reported that almost 90% of forbs in a mesic grassland stocked with cattle at a high density for a short period experienced partial or complete defoliation, tearing, or shredding of their stems and leaves. Abundant forbs have the greatest chance of being damaged, but some rare species also do not escape herbivory (Chamane et al., 2017b). Plant architecture appears to be an important determinant of what direct disturbance forb experience and how they respond to tissue damage by grazing and trampling. Forbs that are reduced or eliminated by chronic herbivory tend to have an erect growth habit, rendering them most exposed, and thus susceptible, to the actions of herbivores, whereas those that proliferate in overstocked grassland are typically prostrate hemicryptophytes that experience less damage to their above-ground tissues or to their buds that are held close to the ground (Uys, 2006; Cole, 1995; Chamane et al., 2017a; Morris and Scott-Shaw, 2019). Very few indigenous forbs can persist in the persistently overgrazed site, which are dominated mostly by non-native ruderal prostrate forbs and grazing-tolerant, course grasses (O’Connor, 2005; Morris and Scott-Shaw, 2019).

Loss of photosynthetic leaf area through grazing, trampling, or mowing would immediately limit carbon sequestration, retarding total above-ground growth of forbs (Chamane et al, 2019). Supply of photosynthates to USOs in geophytic forbs would also be curtailed, reducing the amount of non-structural carbohydrates (NSCs) available for maintenance, storage (Dafni et al., 1981; Chapin et al., 1990), and spring regrowth (Bayer, 1955; Tolsma, 2002; Werger and Huber, 2006; Martínez-Vilalta et al., 2016). Therefore, the negative effect of frequent defoliation in one season on above-and below-ground productivity could be carried over into the following season (White, 1973; Turner et al., 1993; Briske et al., 1996). Frequent herbivory over many seasons could eventually reduce the size and regenerative capacity of for USOs and their potential for long-term survival in heavily grassed mesic grassland swards (Chamane et al., 2019).

The hypothesis that repeated defoliation of aerial parts of a forb during a season would diminish the size of underground storage organs, resulting in reduced plant performance in the spring of the following season was tested in pot trial using *H*. *hemerocallidea*. It was predicted that frequent summer defoliation would: (1) reduce aerial plant production (leaves, inflorescences, and total phytomass) in the season of application, and (2) decrease corm mass as well as aerial biomass production in the subsequent spring. Restricted light conditions in spring were also imposed on defoliated and undefoliated control plants to force them draw fully on their stored reserves rather than jointly or solely on concurrent photosynthesis (Danckwerts and Gordon, 1990) for spring regrowth as a means of further assessing the effect of defoliation history on plant vigour (Wigley et al., 2021). It was predicted that previously defoliated plants would perform most poorly in the dark when drawing on reserves because of the legacy effect of frequent defoliation on diminishing the store of underground energy reserves available for spring regrowth.

## 2. Methods

### 2.1. Study species

*Hypoxis hemerocallidea* Fisch., C.A.Mey. & Avé-Lall. is a long-lived perennial herbaceous geophyte widespread in mesic grassland, savanna and thicket in south-eastern, eastern, central, and northern South Africa as well as Lesotho, eSwatini, Botswana, Zimbabwe, and Mozambique (Williams et al., 2019; Mofokeng et al, 2020). Hairy, sickle-shaped leaves (110-600 mm in height, 10-15 mm wide) grow in three ranks, emerging from terminal buds on an underground corm (25-70 mm diameter; up to 500 g in weight) along with long stems bearing yellow-shaped flowers (6-16 per stem) that open and close daily (Pooley, 1998; Mofokeng et al., 2018). Leaves emerge first in spring and thereafter synanthously with flowers (Pooley, 1998; Lamont and Downes, 2011).

*Hypoxis hemerocallidea* is still currently abundant and of least conservation concern (Williams et al., 2019). However, extensive loss of grassland through land-use change (Jewitt et al., 2015) will reduce its future range while extensive harvesting of the corm for a wide range of traditional and commercial medicinal purposes (Khan and Drewes, 2004; Owira and Ojewole, 2009; Matyanga et al., 2020) depletes local populations, especially around urban centres (Dold and Cocks, 2002; Mofekeng et al., 2020). Plants with large corms are becoming less available in the wild because of overharvesting (Williams et al., 2007).

### 2.2. Study site and experimental design

The experiment was conducted, under light hail netting, at the N.M. Tainton arboretum on the Pietermaritzburg campus of the University of KwaZulu-Natal, South Africa (mean annual rainfall = 844 mm; mean daily temperature in hottest months = 28.2 °C; mean daily minimum winter temperature in coolest months = 2.9 °C). *Hypoxis hemerocallidea* plants obtained from a commercial nursery were grown in 14 litre pots in an organic (> 6% C; 0.55% N), slightly acidic (pH = 5.53) nutrient-rich growing medium (P = 81 mg l^-1^, K = 156 mg l^-1^, Ca = 2960 mg l^-1^, Mg = 410 mg l^-1;^; Manson et al., 2020) in order not to limit their growth. During the summer growing season (2019-2020), pots were watered to saturation on two occasions when no rainfall was received during hot periods. Because rainfall was low (< 20 mm) during the following spring (2020), all plants were given addition water at the end of winter (to initiate regrowth) and weekly in September.

The experiment comprised a full 2 x 2 factorial pot trial arranged in a completely randomised design with five replicate plants of *H*. *hemerocallidea* per treatment combination (n = 20). The treatment factors were frequent defoliation (clipped, unclipped) and spring sunlight growing conditions (dark, full light).

Defoliation was implemented during the first growing season (2019-2020) by clipping all above ground material at a height of 80 mm above the soil surface to remove most of the photosynthetic leaf surface, resulting in a mean defoliation intensity of 84% (77.8-87.1%) reduction in the height of the tallest leaf. Plants were clipped six times, at a mean defoliation interval of 30 days, with shorter between clipping in the summer growing season (17-25 days) and longer intervals in autumn (60 days). The first defoliation was at the beginning of December 2019 and the sixth defoliation was in June 2020. Total above-ground phytomass was harvested at ground level towards the end of winter (23 July 2020) to assess the effect of defoliation on plant performance during the first season.

The carry-over effect of defoliation was assessed by measuring regrowth during the following spring (August – October 2020) with plants growing in full sunlight or under ventilated light-proof boxes that restrict light but allow air circulation for plant growth (Edwards, 1965). Plants regrowing in the dark have to rely on stored energy reserves, i.e. corms in the case of *H*. *hemerocallidea*, whereas those in full sunlight also sequester additional carbon via concurrent photosynthesis; a comparison of their relative vigour indicates the extent to which reserves are available or have been depleted by defoliation during the previous growing season (Muzzell and Booysen, 1995; Peddie et al., 1995; Ripley et al., 2015).

### 2.3. Growth performance measures

Total plant above-ground dry matter (AGDM) yield at the end of the first growing season was measured as the sum of all material removed by each clipping and the final standing phytomass (leaves plus inflorescences) in winter. At the end of spring (6 October 2020) all above-ground phytomass was harvested to ground level and corms were excavated. Roots could not be separated from the polystyrene material used at the bottom of pots for drainage and were therefore not included in below-ground biomass measures. All phytomass was dried to constant mass in an oven at 60 °C. Additional plant measures taken before each clipping, at the end of the growing season (in June 2020), and in the following spring (October 2020) were: number of leaves, number of inflorescences, and height of the tallest leaf, which was typically marginally longer than the other leaves on a plant.

#### Statistical analyses

The effect of defoliation (clipped vs. control) on plant performance (number of leaves, number of inflorescences, leaf height, and AGDM) measured at the end of the first growing season was tested with t-tests for unequal variance. Two-way analysis of variance (ANOVA) was used to assess the main and interaction effects of previous season defoliation (unclipped vs clipped) and spring regrowth light conditions (restricted vs. full light) on the above-mentioned plant measures and dry corm mass. Post-hoc comparisons across factor means were conducted only when interaction effects were significant (p ≤ 0.05). Linear regression was used to assess the relation between AGDM, number of leaves, number of inflorescences, and corm dry mass at the end of spring.

## 3. Results

### 3.1 Treatment season responses

Unclipped control plants grew steadily from an initial maximum leaf height of 48.4 ± 5.15 cm to reach a peak height of 66.0 ± 2.02 cm in early February, diminishing somewhat thereafter to 61.2 ± 1.70 cm at the end of the growing season in early June (Fig. 1). Clipped plants were as tall as control plants at the start (53.8 ± 2.16 cm) and the end (65.0 ± 2.74 cm) of the measuring period in early December and early June, respectively. Clipped plants regrew to a height of 36.0 ± 0.87 cm (63.8% of control plants) after the first defoliation and between 50.0-51.6 cm (75.8-82.2% of control plant height) after the second, third, and fourth clipping (Fig. 1). The mean clipping intensity at a fixed height of 8 cm above soil level was 13.5 % (12.1-16.5%) of control plant heights measured at the same times. At the end of the growing season, control plants had a mean of 20.5 ± 2.32 % dry leaves (> 50% of a leaf brown) but all leaves on all plants had completely senesced by the final harvest date towards the end of July. In contrast, all leaves of all clipped plants remained completely green until June and five plants still retained green leaves in July.

**FIGURE 1:**
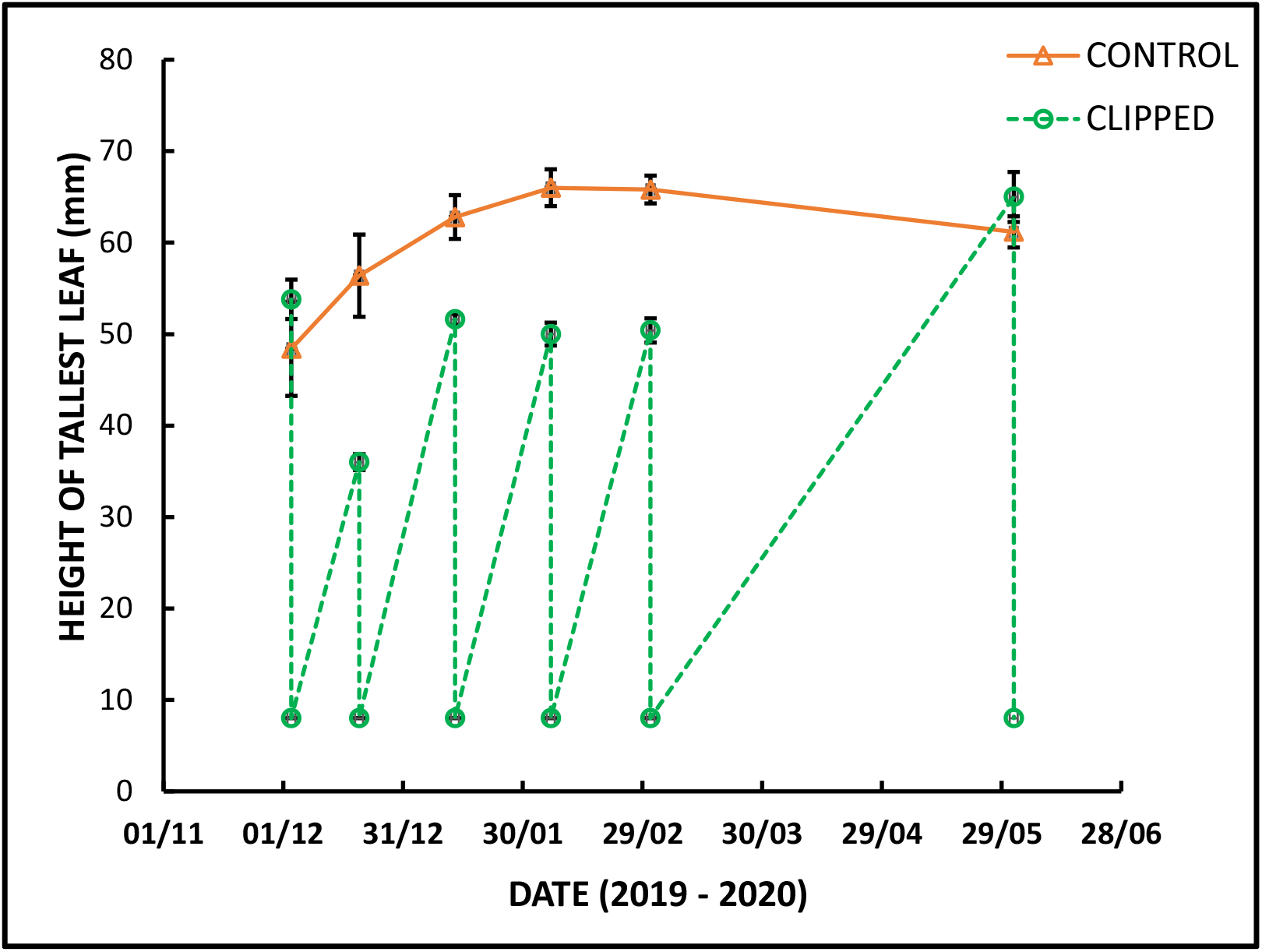
Seasonal changes in the mean (± se) height of the tallest leaf of clipped and unclipped plants of *Hypoxis hemerocallidea*.

Although clipped plants regrew to the same height (t = 0.493, p = 0.630) as controls after the final defoliation, they had 16.3 (−59.7%) fewer leaves (t = 7.043, p < 0.0001) and 15 (−84.3%) fewer flowers (t = 6.061, p = 0.0001), on average, than unclipped plants at the end of the growing season (Table 1). Clipping did not reduce the total yield (offcuts plus final harvest) of above-ground dry matter (AGDM) compared to the control (t = 0.147, p = 0.885) plants (Table 1).

**TABLE 1:** The effect of frequent, severe clipping on plant growth parameters and yield of *Hypoxis hemerocallidea* at the end of the growing season. Significant t-values (p ≤ 0.05) are in bold.

| Variable | CLIPPED |  | CONTROL |  | t-value | p-value |
| --- | --- | --- | --- | --- | --- | --- |
|  | Mean | se | Mean | se |  |  |
| Number of leaves | 11.0 | 0.77 | 27.3 | 2.18 | 7.043 | **0.00002** |
| Number of inflorescences | 2.8 | 0.63 | 17.8 | 2.39 | 6.061 | **0.00012** |
| Maximum leaf height (mm) | 63.7 | 2.89 | 62.1 | 1.48 | 0.493 | 0.63042 |
| Total above-ground dry matter yield (g) | 35.68 | 3.99 | 36.52 | 4.12 | 0.147 | 0.88510 |

### 3.2 Spring regrowth responses

No interactive effects (p > 0.05) of previous-season defoliation and light availability on spring regrowth were evident for any of the plant parameters measured in early October (Table 2). Light regime during spring did, however, have a consistent effect, irrespective of clipping history, on maximum leaf height and inflorescence number with leaves being etiolated by 38% (13.1 mm; p = 0.0004) and plants having 22.4 % (−1.5) fewer inflorescences (p = 0.0246) in the dark than in full sunlight (Table 2; Fig. 2). Restricted light conditions tended to reduce leaf number, but not significantly so (p = 0.0907), but did not affect (p > 0.50) AGDM nor corm dry mass (Table 2; Fig. 2).

**TABLE 2:** Analysis of variance of the effects of clipping (clipped vs. unclipped) and light availability (dark vs. full sunlight) on the regrowth and yield of *Hypoxis hemerocallidea* in October measured at the end of the spring growth period. Significant t-values (p ≤ 0.05) are in bold.

| Effect | d.f. | No. leaves |  | No. inflorescences |  | Max. leaf height |  | AGDM<sup>1</sup> yield |  | Corm dry mass |  |
| --- | --- | --- | --- | --- | --- | --- | --- | --- | --- | --- | --- |
|  |  | F | p | F | p | F | p | F | p | F | p |
| Light | 1 | 3.24 | 0.0907 | 6.16 | **0.0246** | 19.91 | **0.0004** | 0.37 | 0.5515 | 0.36 | 0.5569 |
| Defoliation | 1 | 1.11 | 0.3077 | 23.04 | **0.0002** | 0.42 | 0.5261 | 10.54 | **0.0051** | 14.34 | **0.0016** |
| Light x Defoliation | 1 | 0.14 | 0.7132 | 1.34 | 0.2640 | 0.14 | 0.7132 | 0.70 | 0.4151 | 0.000039 | 1 |
| <sup>1</sup>AGDM = above-ground dry matter. |  |  |  |  |  |  |  |  |  |  |  |

**FIGURE 2:**
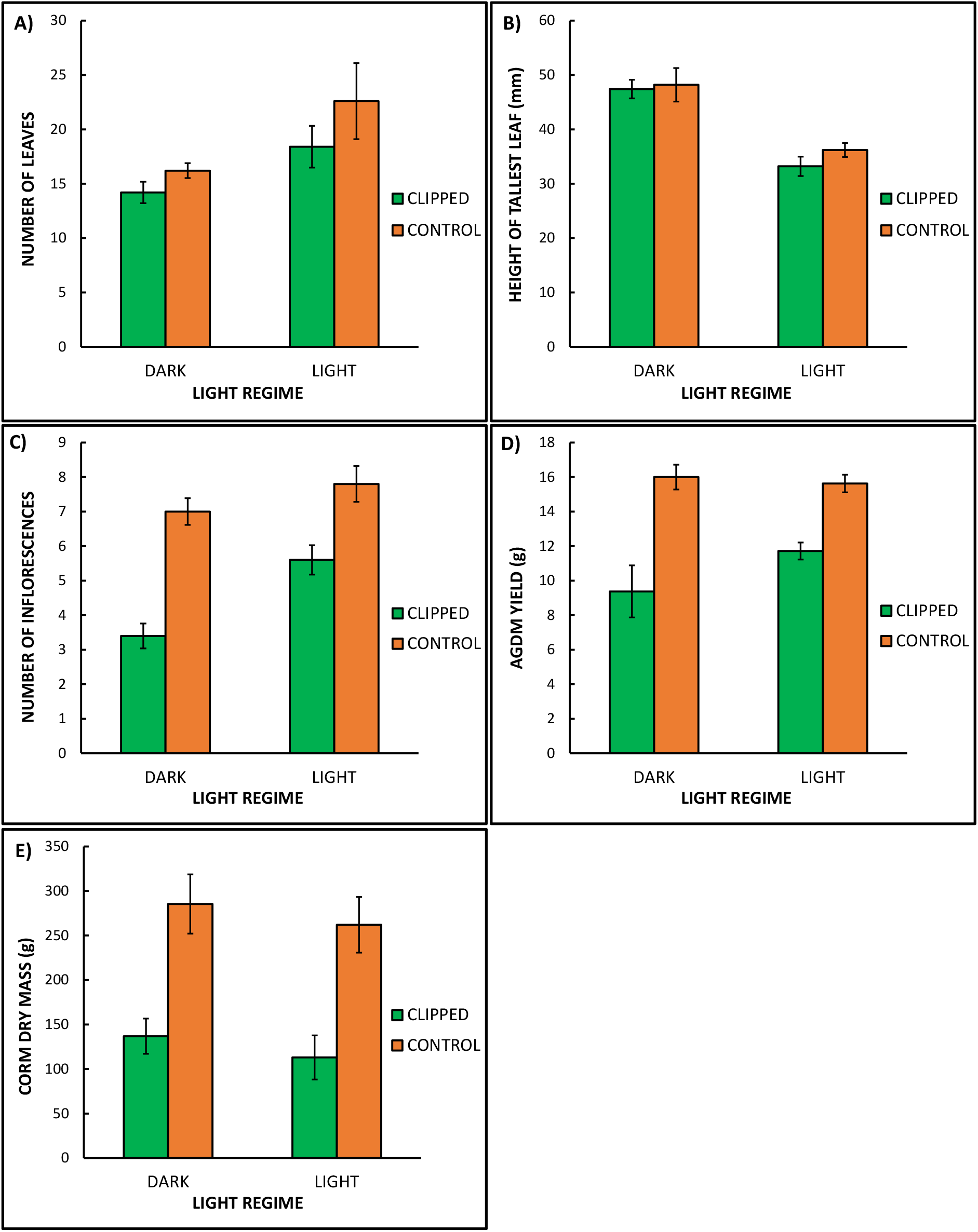
Mean (± se) number of leaves (a), height of the tallest leaf (b), number of inflorescences (c), above-ground dry matter (AGDM) yield (d), and corm mass (e) of clipped and unclipped (control) plants of *Hypoxis hemerocallidea* at the end of spring in October.

Clipping five times during the preceding growing season consistently reduced inflorescence number (−2.9) by 39.2% (p = 0.0002) compared to undefoliated controls but did not influence the number of leaves (p = 0.308) nor their maximum height (p = 0.526) at the end of spring (Table 2; Fig. 2). Above-ground yield was reduced by one-third (−5.27 g; p = 0.0051) and corm mass was more than halved (−54.4%; - 148.4 g; p = 0.0016) by frequent clipping during the previous season (Table 2; Fig. 2).

At the end of spring, there was a positive correlation between AGDM yield and corm mass of *H*. *hemerocallidea* plants (r =0.552, p = 0.012). Above-ground dry matter varied the most for plants with small corms (Fig. 3). Number of inflorescences (r = 0.461, p = 0.041), but not leaf number (r = 0.139; p = 0.559), was positively correlated with corm dry mass.

**FIGURE 3:**
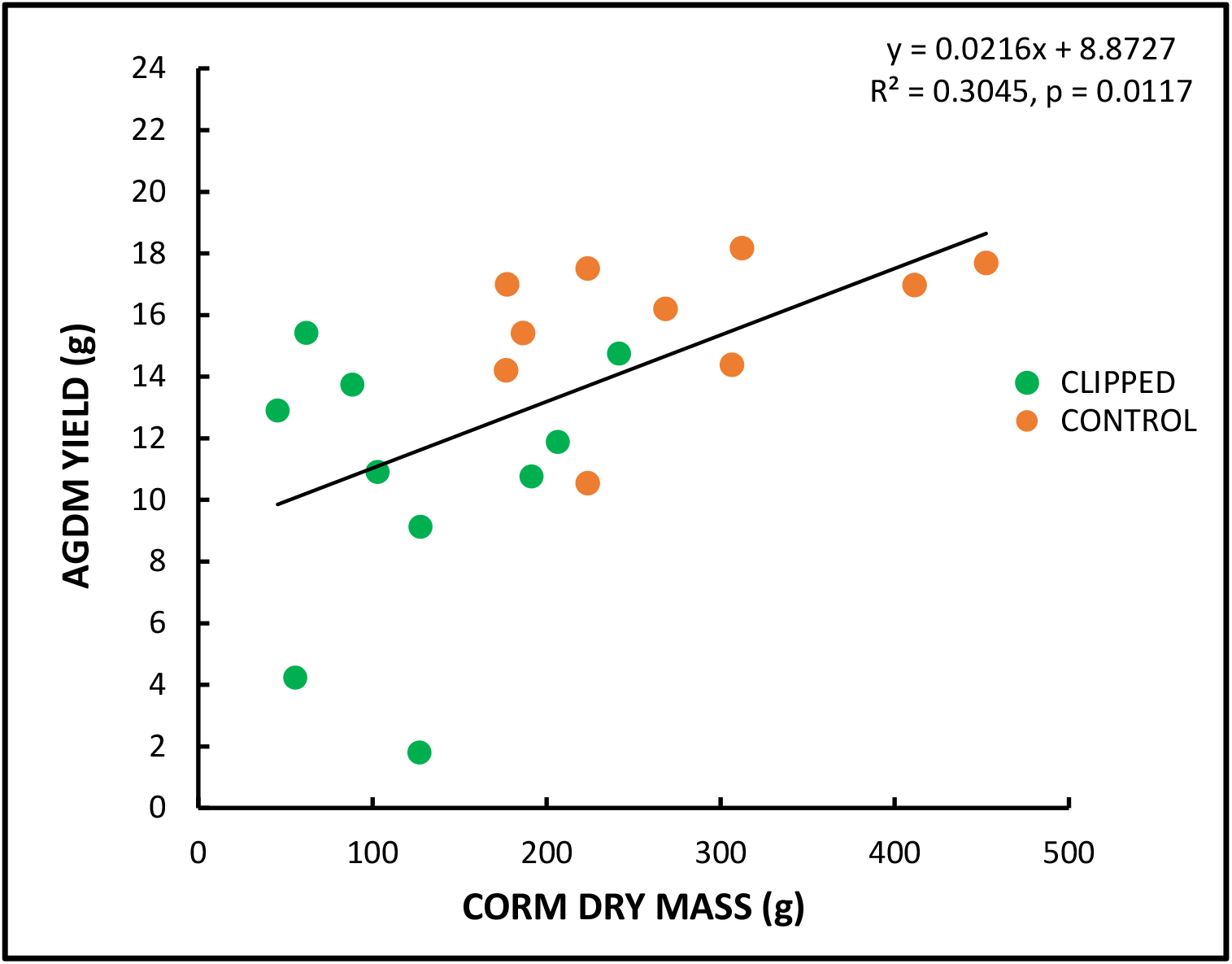
Linear regression of above-ground dry matter (AGDM) yield on corm dry mass of clipped and unclipped (control) plants of *Hypoxis hemerocallidea* at the end of the spring in October.

## 4. Discussion

This study set out to explore a possible mechanism to explain the demise of many mesic grassland forbs in overgrazed mesic grassland by examining the immediate and seasonal carry-over effects of recurrent defoliation on the above- and below-ground performance of a mesic grassland geophytic forb, *H*. *hemerocallidea*. It was hypothesised that frequent leaf damage by herbivory, simulated by severe clipping, would reduce above-ground plant growth and that the corms (the USO) of the *H*. *hemerocallidea* would be diminished by recurrent defoliation because of reduced carbon supply from leaves for storage and a greater demand for stored reserves for regrowth. Key results in support of this hypothesis were that plants were largely resilient to repeated, severe defoliation in the first season of cutting but regrowth was reduced in the following spring. Corm mass was more than halved by defoliation, indicating that USOs ostensibly protected underground are vulnerable to adverse effect of disturbance to aerial plant parts. Herbivore induced depletion of USOs could, therefore, potentially threaten the persistence of forbs in grassland chronically disturbed by heavy grazing or frequent mowing.

The importance of USOs for supplying energy and other resources to forb species that need to resprout after a dormant period was supported by the results of this experiment. *Hypoxis hemerocallidea* plants that grew under restricted light conditions, when they had to fully rely on stored energy reserves rather than on concurrent photosynthesis for spring leaf production, were able to match the total growth of plants that resprouted in full light, although their flower production was somewhat curtailed (−22.4%). Forbs that have access to a substantial buried bud bank (Klimešová and Klimes, 2007) and a store of resources can initiate growth sooner than grasses, thereby exploiting the limited opportunity to sequester carbon, grow, spread, and produce flowers, before light becomes limiting because of grass canopy closure (Bayer, 1955; Lammont and Downes, 2011). Bews and Vanderplank (1930) found that such spring growth does not appear to significantly deplete NSCs (primarily starch) in the corms of *H*. *hemerocallidea* dug up periodically from the field and questioned the value of stored energy reserves for regrowth. They did, however, note that other resources such as minerals and water (Chapin et al., 1990; da Silva and Rossatto, 2019) are also supplied by the USO (Bews and Vanderplank, 1930). Plants may continue to draw up nitrogen and phosphorus into leaves after they have ceased mobilising NSCs from the USO to initiate regrowth (Rosenthal and Kotanen, 1994). Buried storage organs also enable grassland forbs to escape frost, tolerate temporary drought during the dry autumn to winter period, and survive fires that occur commonly during or shortly after the dormant period (Dafni et al. 1981; Ruiters et al., 1993; Proches et al. 2005; Clarke et al. 2013). Directing growth to protected storage organs, does therefore, appear to be a key plant survival trait for environments where there is a high, persistent risk of complete loss of leaves through senescence or fire during the dry season (Ott et al., 2019; Siebert et al., 2019; Ottaviani et al., 2020).

Underground stores also enable plants to tolerate partial or complete damage or loss of leaves by mowing, insect folivory, or mammalian herbivory when they are actively growing (Strauss and Agrawal, 1999; Tiffin, 2000; Thomas et al., 2017). The substantial corm of *H*. *hemerocallidea* likely enabled it to fully compensate for biomass lost through frequent, severe defoliation during the first summer growing season by providing resources for repeated regrowth. Such a compensatory growth response indicates that, although the imposed defoliation regime would have limited carbon sequestration, the overall availability of carbon to the plant for repeated regrowth was not limiting (Wise and Abrahamson, 2005). Leaf and flower numbers, were, however, significantly impacted by defoliation, possibly because of a prioritisation of leaf extension and expansion over initiation of new leaf and floral primordia in the corm (Shefferson et al., 2006).

The effect of growing-season disturbance intensity, frequency, and timing on biomass allocation and growth of geophytic forbs has, unlike for grasses (Briske and Richards, 1995; Ferraro and Oesterheld, 2002), received very little attention. Frequent mowing suppresses above- and below-ground forb biomass (Ottaviani et al., 2021) and forb species diversity (Fynn et al., 2014). Generally, longer periods between defoliations (i.e., low defoliation frequency) reduce the negative effect of defoliation on plants (Ferraro and Oesterheld, 2002) by allowing leaf area to accumulate, roots to grow and stores to be replenished and accumulated (Bayer, 1955; Chapin et al., 1990; Tyler and Borchert, 2003). The latter part of the growing season from late summer into autumn, when growth rates slow under declining temperatures but photosynthesis continues, could be a crucial period for storage (Chapin et al., 1990) and recycling of nutrients from senescing leaves down to USOs (Ruiters et al., 1993). The seasonal carbohydrate and nutrient dynamics of *H*. *hemerocallidea* and other mesic grassland geophytes in relation to disturbance have not been examined but affording plants an opportunity to recuperate and accumulate excess non-structural carbohydrate reserves, especially starch, in autumn could enhance their ability to rapidly resprout in the following spring (Janeček et al., 2015; Martínez-Vilalta et al., 2016). Clipped plants of *H*. *hemerocallidea* retained a higher proportion of green leaf than unclipped plants during autumn, perhaps contributing to their ability to recover their stature and biomass production after frequent defoliations in summer. This resilience displayed by *H*. *hemerocallidea* under summer defoliation was, however, not repeated in the following spring when previously clipped plants produced a third-less above-ground biomass than unlipped plants.

The most marked impact of single season of clipping on *H*. *hemerocallidea* was the loss of more than half of its corm mass. Ottaviani et al. (2021) also observed far lower (by > 70%) rhizome biomass in temperate grassland mown twice versus grassland mown once annually. In that study, below- and above-ground biomass scaled linearly (Ottaviani et al. (2021), similar to the positive linear relation between corm size and spring biomass production in *H*. *hemerocallidea* (Fig. 3). Lubbe et al. (2021) also found a significant positive correlation between below-ground stored carbohydrate concentration and leaf economic traits, particularly plant height, of 78 temperate herbs. These results on the positive influence of USO size on growth accord with widespread knowledge in the horticultural industry that plants with large bulbs or tubers produce are the most vigorous, producing the highest number of leaves, flowers, and clonal offspring (e.g. Rees, 1969; Ahmad et al., 2009; Kapczyńska, 2014). Tuber and bulb size or mass is a general indication of the total carbon pool available for growth and reproduction (Werger and Huber, 2006; Lee et al., 2016). Therefore, a large reduction in the mass of USOs of a grassland forb resulting from frequent reduction of photosynthetic surfaces area during the growing season, as observed in this trial, would reduce the plant’s ability to recuperate and its long-term resilience to recurrent leaf disturbance (Canadell and López-Soria, 1998; Tolsma, 2002).

Reproductive buds are also borne on the USO of geophytes (Klimešová and Klimes, 2007; Pausas et al., 2018), and any diminution of individual storage organs by grazing (Qian et al., 2017) could reduce the potential for forbs to reproduce sexually or to spread through clonal growth (Klimešová and Klimes 2007). Inflorescence production was the growth component of *H*. *hemerocallidea* most markedly reduced by clipping, both during the season of clipping and in the following spring; smaller corms bore significantly fewer inflorescences. This suggests that initiation and extension of floral buds on the corm was not prioritised in the allocation of resources mobilised from the USO for regrowth (Chapin et al., 1990). Seed production in *H*. *hemerocallidea* is high but germination is inherently poor because of a hard seed coat and embryo dormancy (Hammerton and van Staden, 1988; Katerere, 2015).

Seedling emergence and successful recruitment by mesic grasslands forbs is generally very low (Uys, 2006) and seedlings could be vulnerable to the grazing and trampling of herbivores (Scott-Shaw and Morris, 2006; Chamane et al., 2017b). Consequently, the already tight demographic bottleneck of forbs could be narrowed by herbivory, reducing the likelihood of new recruits. *Hypoxis hemerocallidea* plants can produce cormlets from corm buds, with medium to large corms potentially producing the most daughter corms (Mofokeng et al., 2018). However, adults have not been noted to occur in clumps (Pooley, 1998), which suggests that clonal expansion of populations is not typical.

Taken together, results of this trial indicate that growing-season damage to leaves, such as by intensive mowing or by the grazing and/or trampling of livestock can reduce the short-term resilience, and potentially long-term persistence of, geophytic forbs such as *H*. *hemerocallidea* in disturbed mesic grassland. Use of stored reserves for regrowth after repeated, intense disturbance during the growing season depletes USOs while diminished USOs reduce the potential for future shoot growth and reserve replenishment (Ottaviani et al., 2021). This recursive, negative feedback on plant growth may result in progressive run-down of vigour and above- and below-ground plant biomass if grazing and trampling continues unabated (Turner et al., 1993). Erect forbs would be most vulnerable to long-term recurrent leaf damage or loss (Fynn et al., 2004; Chamane et al., 2017a; Morris and Scott-Shaw, 2019). Gradual attrition of USOs and bud banks under chronic herbivory could, therefore, be the mechanism whereby populations of mesic grassland forbs are extirpated or severely reduced through overgrazing (Scott-Shaw and Morris, 2015).

This defoliation pot experiment on *H*. *hemerocallidea* has provided proof of the principle that recurrent growing-season disturbance to aerial parts of a geophytic forb negatively impacts buried storage organs and spring recovery growth, with a potential impact on its longevity in highly disturbed environments. The application of this principle to other geophytic forb species in a variety of environments would depend on several factors. In grazed fields, forbs are likely to experience less frequent and intense defoliation – with perhaps only partial leaf damage - than the experimental levels imposed upon *H*. *hemerocallidea*. Therefore, the short-term impact of herbivores on shoots and USOs would probably be smaller than that observed in this trial. However, *H*. *hemerocallidea* has a substantial corm that might better tolerate frequent defoliation than the smaller USOs of those mesic grassland forbs that resprout from woody rootstocks, thickened (e.g., fusiform) roots, or rhizomes (Pooley, 1998; Uys, 2006); such species could thus be especially vulnerable to herbivory. The capacity for compensatory growth is generally highest for plants growing in fertile soils (Wise and Abrahamson, 2005), such as that used in this trial where nutrients were not limiting. On less fertile soils and in drier, less productive environments, plants would be less tolerant of recurrent loss of tissue (Janeček et al, 2015; Morris, 2016) and, consequently, more likely to be severely impacted by sustained disturbance. Further research is thus required on the seasonal and carry-over effects of leaf disturbance on forbs with a range of USO types and sizes, their long-term response to herbivory, and management options for sustaining the abundance and diversity of forbs in mesic grassland.

Careful management of extant populations of forbs in mesic grassland is required because once eliminated from primary grassland, such long-lived perennials do not readily reestablish (Zaloumis and Bond, 2016; Buisson et al., 2019) even after many centuries of recovery (Nerlekar et al., 2020). Three key management interventions are required to maintain forbs in grazed mesic grassland: regular fire, lenient grazing or mowing, and periodic rests. Without regular disturbance by burning during the dormant season or early spring, forb diversity, composition, and underground bud banks will decline (Uys et al., 2004; Fynn et al., 2005; Fidelis et al., 2014; Kirkman et al., 2014; Gordijn et al., 2018). Forb biomass and diversity can also be maintained by infrequent summer mowing (Fynn et al., 2004; Ottaviani et al., 2021). Heavy stocking with concentrated herds of livestock is not a safe alternative to fire or mowing, as advocated by proponents of high-density grazing systems (e.g. Savory and Butterfield, 2016) because hooves repeatedly damage living leaves (Chamane et al., 2017b), eventually leading to the demise of many forbs (Scott-Shaw and Morris, 2015). In contrast, moderate livestock stocking rates and densities, especially when coupled with controlled burning (O’Connor et al., 2010; SANBI, 2014; Scott-Shaw and Morris, 2015; Joubert et al., 2017) are sustainable without the risk of damaging forbs and ecosystem processes. However, even lightly stocked mesic grassland needs to be afforded a full-season rest (deferment) from grazing every few years to restore the vigour of grazed palatable grasses (Kirkman, 2002; McDonald et al., 2019). Such periods of uninterrupted growth would likely also benefit forbs which have experienced extensive leaf damage and USO depletion over several consecutive years by allowing them to replenish their reservoir of stored reserves (Janeček et al., 2015; Martínez-Vilalta et al., 2016) and to flower and set seed (Uys, 2016).

## 5. Conclusion

Mesic grassland forbs are fire-adapted but not grazing resistant. Their underground storage organs (USOs) protect them from frost and regular dormant-season fires, but, as this research demonstrates, below-ground plant organs are sensitive to above-ground disturbance in the growing season. Recurrent damage to growing leaves can negatively impact forbs, markedly diminishing the biomass of their USO and their consequent ability to support resprouting in spring. Therefore, the regenerative and resource storage organs of long-lived geophytic forbs located underground are not safe from excessive mowing or herbivory (cf. Klimešová et al., 2018). The observed marked negative carry-over effect of frequent defoliation of *H*. *hemerocallidea* on its subsequent regrowth and corm size portends dwindling resilience and potentially lower longevity of plants under unrelenting intense disturbance. A substantial reduction in flowering in perennial mesic grassland geophytic herbs could be a useful bellwether of declining vigour in forbs because inflorescence production is highly sensitive to herbivory.

Management of species-rich mesic grassland needs to aim to minimise repeated, severe damage to the green above-ground biomass of forbs. Management during the growing season should, therefore, be lenient and high stocking densities should be avoided. Infrequent summer mowing or grazing at moderate stocking rates and densities combined with fire and periodic full season resting are recommended to maintain the vigour and resilience of forbs in mesic grassland.

## 6. Acknowledgements

I am grateful to Welcome Ngcobo who helped set up and maintain the experiment, Anita Morris for her help in treatment application and data collection, and Nicky Findlay for the soil analysis.

